# Transcriptome landscape of kleptoplastic sea slug *Elysia viridis*

**DOI:** 10.1101/2022.06.28.497858

**Authors:** Manuel Mendoza, Sara Rocha, Jesús Troncoso, David Posada, Carlos A. Canchaya

## Abstract

Certain sacoglossan sea slugs can sequester and maintain photosynthetically active chloroplasts through algae feeding, a phenomenon called kleptoplasty. The period while these plastids remain active inside the slug’s body is species- and environment-dependent and can span from a few days to more than three weeks. Here we report for the first time the transcriptome of sea slug Elysia viridis (Montagu, 1804), which can maintain kleptoplasts for more than two weeks and is distributed along all the Atlantic European coastline. The obtained transcriptome of E. viridis comprised 12,884 protein-coding sequences (CDS). The shortest one was 261bp, and the longest 8,766bp; the whole transcriptome has a total length of 9.3Mb (Table S4 and Fig. S2). Analysing these CDS, we identified 9,422 different proteins, with best hits mainly from two genera: Elysia (87.2%), and Plakobranchus (11.0%) (Fig. S2); the other 2.3% corresponded to multiple genera of sea slugs and snails (Tectipleura) (Kano et al., 2016). We got the functional annotation (Gene Ontologies, GO) corresponding to 9,333 CDS: 4,755 CDS associated with 2,583 Biological Process (BP); 5,466 CDS linked to 683 Cellular Components (CC); and 6,693 related to 1,606 Molecular Functions (MF). We identified 201 CDS related to response to stress (GO:0006950) and 10 CDS associated with the regulation of response to stress (GO:0080134). Focussing on the ROS-quenching toolkit, we found 24 CDS related to oxidoreductase complex (GO:1990204) and 560 annotated with oxidoreductase activity (GO:0016491) acting in a large number of donors, e.g., CH-OH, CH=O, C=O, CH and CH2. In addition, we found 39 CDS with antioxidant activity (GO:00162099) and other CDS with ROS-quenching function: superoxide dismutase (GO:0004784), peroxidase (GO:0004601), glutathione oxidoreductase (GO:0097573) and peroxidase (GO:0004602); and thioredoxin peroxidase (GO:0008379) activity. Furthermore, we found 8 CDS related to the symbiont response (GO:0140546) and nine related to the pattern recognition receptor signalling pathway (GO:0002221).

## RESEARCH NOTE

Certain sacoglossan sea slugs sequester photosynthetic-active chloroplasts of their prey, recognising them through the pattern recognition receptors (Melo Clavijo *et al*., 2020). The hosts rearrange their transcriptional landscape (Chan *et al*., 2018) and keep the kleptoplasts inside the epithelial cells of digestive tubules (de Vries *et al*., 2014). Here, the stolen plastids play a dual role as starch-storage device and nutritive source during scarcity periods (Cartaxana *et al*., 2017; Laetz *et al*., 2017), even though their photosynthates are not essential for the slug’s nutrition during starvation (Christa, Zimorski, *et al*., 2014). The time while kleptoplasts remain functional and active is different among the multiple species, from a few days (short-term retainers, StR) to over one month (long-term retainers, LtR) (Händeler *et al*., 2009).

From the beginning, the kleptoplasts are challenged by the host’s innate immune system, the first line of defence against potential pathogens. The immune response of molluscs is triggered by recognising evolutionary conserved pathogen-associated molecules through the different types of pattern-recognition receptors (PRR) (Cao, 2016). One group of these, the scavenger receptors, may play a crucial role at the beginning of kleptoplasty (Melo Clavijo *et al*., 2020). After chloroplast recognition, the host rearranges its transcriptional landscape, increasing the expression of certain genes, e.g. reactive oxygen species (ROS) quenching genes (Chan *et al*., 2018).

During starvation periods, damaged photosynthesis-related proteins can generate abundant ROS (Maeda *et al*., 2021), that can oxidise other proteins or organelles, triggering an autophagy signal to reduce the oxidative damage (Scherz-Shouval, Elazar, 2011). Different organisms have also developed multiple mechanisms to neutralise the cytotoxicity of ROS, such as enzymatic reactions and synthesis of other detoxifying agents (Birben *et al*., 2012). Based on the ROS-quenching response during starvation, de Vries *et al*., 2014 proposed a new classification for the photosynthetic sea slugs, i.e., starvation-intolerant and starvation-tolerant species, depending on the ability to activate the expression of a quenching burst to control the proliferation of ROS.

To shed more light on mechanisms behind the kleptoplasty, we describe here for the first time the transcriptomic landscape of the photosynthetic sea slug *Elysia viridis* (Montagu, 1804), by assembling de novo a reference transcriptome from a pool of ten individuals. This species is a facultative non-selective Ulvophyceae-feeder LtR sacoglossan sea slug found across the European Atlantic border, from Scandinavia to the British Isles and the Iberian Peninsula (Jensen, 2007). *E. viridis* has a wide range of kleptoplasts’ retention times depending on the plastid source (Christa, Händeler, Schäberle, *et al*., 2014; Rauch *et al*., 2018). We share the reads and the assembled transcriptome through the NCBI BioProject accession number PRJNA549923. We focused our description on two points: (1) characterisation of different types of pattern recognition receptors (PRR), focusing on the thrombospondin type 1 repeat (TSR) superfamily, scavenger receptors (SR), and Toll-like receptors (TLR); and (2) identification and characterisation of orthologs related to kleptoplasts retention. We followed the common pipeline for transcriptome *de novo* assembly and annotation (Conesa *et al*., 2016; Raghavan *et al*., 2022). We described in detail the pipeline used in the following repository: manuelsmendoza/elvira.

The obtained transcriptome of *E. viridis* comprised 12,884 protein-coding sequences (CDS). The shortest one was 261bp, and the longest 8,766bp; the whole transcriptome has a total length of 9.3Mb (Table S4 and Fig. S2). Analysing these CDS, we identified 9,422 different proteins, with best hits mainly from two genera: *Elysia* (87.2%), and *Plakobranchus* (11.0%) (Fig. S2); the other 2.3% corresponded to multiple genera of sea slugs and snails (Tectipleura) (Kano et al., 2016). We got the functional annotation (Gene Ontologies, GO) corresponding to 9,333 CDS: 4,755 CDS associated with 2,583 Biological Process (BP); 5,466 CDS linked to 683 Cellular Components (CC); and 6,693 related to 1,606 Molecular Functions (MF).

We identified 201 CDS related to response to stress (GO:0006950) and 10 CDS associated with the regulation of response to stress (GO:0080134). Focussing on the ROS-quenching toolkit, we found 24 CDS related to oxidoreductase complex (GO:1990204) and 560 annotated with oxidoreductase activity (GO:0016491) acting in a large number of donors, e.g., CH-OH, CH=O, C=O, CH and CH_2_. In addition, we found 39 CDS with antioxidant activity (GO:00162099) and other CDS with ROS-quenching function: superoxide dismutase (GO:0004784), peroxidase (GO:0004601), glutathione oxidoreductase (GO:0097573) and peroxidase (GO:0004602); and thioredoxin peroxidase (GO:0008379) activity. Furthermore, we found 8 CDS related to the symbiont response (GO:0140546) and nine related to the pattern recognition receptor signalling pathway (GO:0002221).

Melo Clavijo *et al*., 2020 proposed a recognition of kleptoplasts by the SRs and C-type lectin receptors (CTLR) mediated by thrombospondin type 1 repeat (TSR) protein superfamily, similar to the theory proposed by (Neubauer *et al*., 2017) describing the recognition of algae symbionts by cnidaria. In our analysis, we identified 19 CDS from the TSR superfamily (Fig. S3) and multiple PRRs (Li, Wu, 2021) (Fig. S3 to S7) that we can classify in three membrane-bound receptors that may be involved in plastid recognition: TLR4, SR-E and CTLR-6. These receptors were found in other Elysoids too (Melo Clavijo *et al*., 2020), though the structure was different in each specie.

In addition, we also found two CDS as semaphorin 5 (SEMA5), a TSR family characteristic of vertebrates (Goodman *et al*., 1999; Pasterkamp, 2012). However, it has been described in different invertebrates (Fig. S4) (Melo Clavijo *et al*., 2020; Gerdol *et al*., 2020; Maeda *et al*., 2021). Subsequently, a recent hypothesis suggested that the SEMA5 family originated in a common ancestor of Placozoa, Cnidaria and Bilateria, though they are absent in Nematodes (Junqueira Alves *et al*., 2019).

The kleptoplastic phenomenon has multiple independent origins across the tree of life (Christa, Händeler, Kück, *et al*., 2015). However, the different groups of slugs, i.e., long-term and short-term retainers, should have common mechanisms that allow those species to steal the plastids, and those tools should be absent in the non-kleptoplastic species. Thus, we compare the orthogroups found in different organisms, depending on their ability to keep the plastids alive or not, as well as LtR and StR species to find the mechanisms behind this classification. The information about different species used and the analysis pipeline are described in the supplementary material. The information about the species is shown in Table S5, and the assembly statistics of those species are stored in Table S6.

We detected that 16,997 orthogroups made up 96% of total CDS. 311 orthogroups were species-specific, while 1,947 were common to all the species (Fig. 2); 37% of orthogroups contains 1 or more gene per specie (on average). 573 orthogroups were specific to (found in all) kleptoplastic species, 109 were exclusive to the short-term retainers species, and 4 were exclusive to the long-term retainer slugs (*sensu lato*). Analysing the LtR elysoids species attending to the plastid source, 47 orthogroups were specific to the Ulvophyceae-feeder slugs, and 70 were associated with the Heterokonts-feeder species (Table S7).

**Figure 1:**
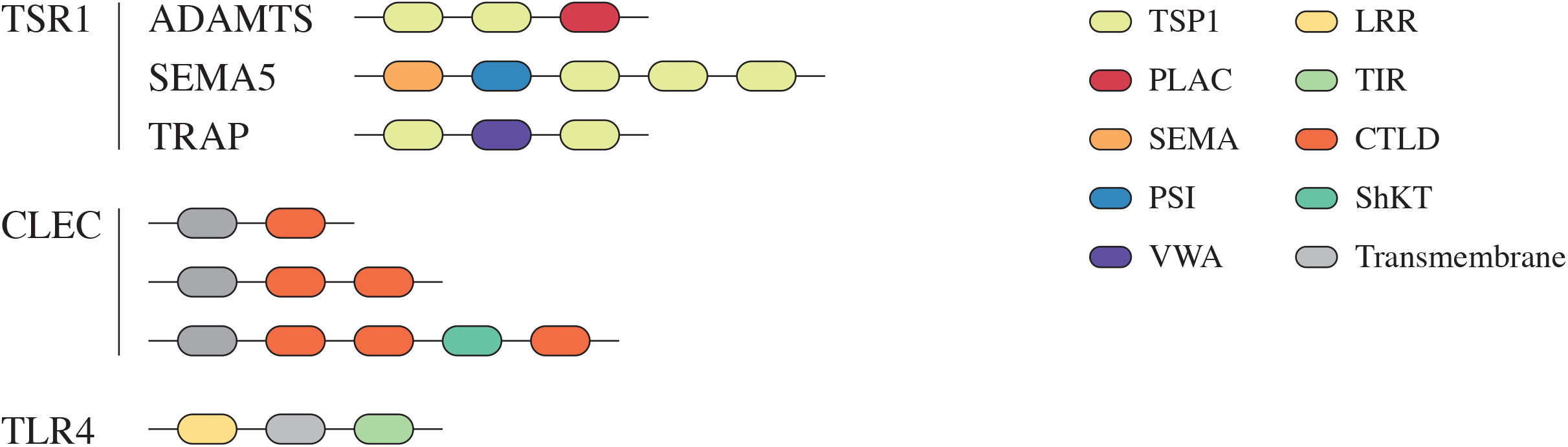
Summary of the PRR found in the transcriptome of *E. Viridis* (All the PRR-coding sequences are described in the supplementary material, Fig. S3 to S7). TSR1 (thrombospondin 1 receptors), ADAMTS (a disintegrin and metalloproteinase with thrombospondin motifs), SEMA5 (semaphorin-5), TRAP (thrombin receptor-activating peptides), CLEC (C-type lectin) and TLR4 (Toll-like receptors 4).

**Figure 2:**
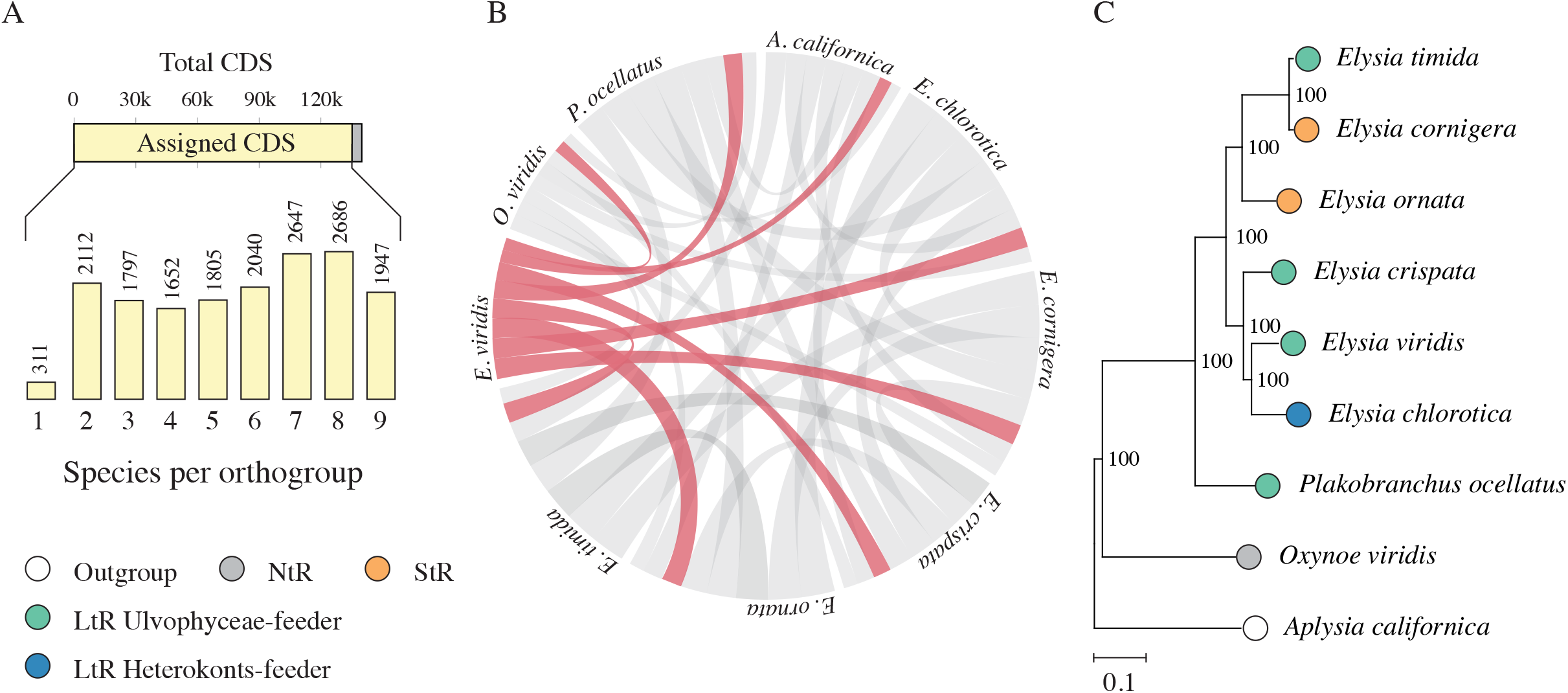
Orthogroups detection and phylogeny. (A) Number of CDS assigned to the different orthogroups and number of species present in each one. (B) Visual representation of the number of orthogroups shared between the different species, in red is the number of orthogroups that share *E. viridis* with the other slugs. (C) A maximum likelihood phylogenetic tree inferred using Rax-ML from 1,380 orthogroups (More info in the Table S8 and Github repo: Phylogenetic analysis); the tree was done under the JTT+FC+I+R4 model and 10,000 bootstrap steps (branches labels show bootstrap percent).

The different kleptoplastic slugs have developed an important burst to regulate the pH, using the iron ions (GO:0006885, GO:0006879, GO:0008199, and GO:0004322) and other enzymatic pathways (GO: 0016491, GO: 0016788 andGO:0016817). Focusing on the LtR species, we found ontologies enrichment related to the immune response (GO:0006955) mediated by the tumour necrosis factor (TNF) (GO:0005164 and GO:0016021) and G-proteins (GO:0004930, GO:0016021 and GO:0007186).

In conclusion, the longevity of kleptoplasts and sea slugs during starvation may be mediated by multiple factors. From the recognition of the plastids during feeding by multiple receptors, including PRR, CTLR and SR and the ROS quenching using enzymatic and non-enzymatic mechanisms. In the particular case of E. viridis we have found the presence of multiple PRR that may be involved in the plastid-recognition process, despite this specie has a lower receptors richesneess in comparison with other Elysoids (Melo Clavijo *et al*., 2020). In addition, we also detected a huge ROS-quenching burst mediated by multiple enzymatic families, whereas the production of antioxidan compounds may have a minor contribution in the control of oxidative stress. Moreover, with a evoluationary perspective, based on the results obtained comparing the different species we could conclue that the platid recognition may require the G-protein-coupled receptors, and the ROS-quenching burst may require Fe-mediaded mechanisms.

## ACKNOWLEDGEMENT

This work was supported by the Consellería de Educación e Ordenación Universitaria, Xunta de Galicia (GPC2014/067) and the Universidade de Vigo to David Posada; and from the European Social Fund and the Government of Xunta de Galicia (ED481A-2018/305) to Manuel Mendoza. We thank the Supercomputing Centre of Galicia (CESGA) for providing computational resources and technical support.

