## Supplementary information for "Transcriptome landscape of kleptoplastic sea slug *Elysia viridis*"

[1] *CIM, Universidade de Vigo, Comparative Genomics Lab., 36310 Vigo, Spain*; [2] *Dept. of Biochemistry, Genetics and Immunology, School of Biology, 36310 Vigo, Spain*; [3] *CINBIO, Universidade de Vigo, Phylogenomics Lab., 36310 Vigo, Spain*; [4] *CIM, Universidade de Vigo, ECOCOST Lab., 36310 Vigo, Spain*; [5] *Dept. of Ecology and Animal Biology, School of Marine Sciences, 36310 Vigo, Spain*; [6] *Galicia Sur Health Research Institute (IIS Galicia Sur), SERGAS-UVIGO, 36213 Vigo, Spain.*

SUPPLEMENTARY INFORMATION

**3 Phylogeny** **12**

### 1 Experiment design and analysis pipeline

Sea slugs from the superorder Sacoglossa can sequester functional chloroplast through feeding and keep them photosynthetically active inside their digestive tubules (de Vries *et al.*, 2014). A small polyphyletic group of sacoglossan species can maintain the stolen plastids (kleptoplasts) functional for more than a month (Händeler *et al.*, 2009). Despite an extensive research record (De Vries *et al.*, 2014), some questions remain unveiled: How are the plastids recognised from the other components of prey's cells? (Melo Clavijo *et al.*, 2020) Why do the sea slugs remain alive after weeks of starvation if the photosynthates are not essential (Christa, Zimorski, *et al.*, 2014)? Even if the ability to sequester the plastids has multiple independent origins along the evolution, can we find orthologs related to the time the plastids remain active? (Christa, Händeler, *et al.*, 2015; Hirokane *et al.*, 2022).

We tried to bring some evidence about the retention time of the different species by sequencing a novel transcriptome from a cosmopolitan species present along the European Atlantic shore, *E. viridis* (K. R. Jensen, 2007) and comparing it with the ones from other species of sea slugs. The sample collection and the methodology used to find the answers are described below.

#### 1.1 Sample collection and maintenance

We sampled wild *E. viridis* individuals from a subtidal *Codium* spp. meadow settled on a granite outcrop (Cabo Estai, Spain, 42°10'5"N 8°48'48"W in October $17^{th}$, 2012). After starving for two weeks in the laboratory under controlled conditions: natural light cycle and sea surface temperature using 0.2$\mu m$-filtered sea water in continuous circulation. We made four pools of ten individuals, preserving them in RNAlater$^{TM}$ at room temperature. Afterwards, we sent the four samples (pools) for RNA extraction, library preparation and Illumina paired-end sequencing to BGI Genomics (China). None of the samples sent passed the quality control required by the technical staff (Table S2). Even so, we ordered to sequence the sample EP3A despite its high fragmentation, it was the single one with enough material, and the RIN might have been underestimated due to the co-migration of both ribosomal fragments (Adema, 2021). The output obtained was a paired-end 90bp length library with 27 million reads.

Table S1: Quality data submitted by the technical staff from BGI. The standard quality requirements for RNA-Seq were: 10g of total RNA and RIN 7. The total RNA mass was measured using a Qubit fluorometer, and the RNA Integrity Number (RIN) was calculated by electrophoresis migration using Agilent Bioanalyzer. For molluscan samples, the RIN should be corrected considering the co-migration of 18S and 28S rRNA in molluscs that may generate false poor-quality results (Adema, 2021).

| Sample name | Total RNA ($\mu$g) | RIN | QC OK? |
|---|---|---|---|
| EP1A | 8.80 | 5.9 | FALSE |
| EP2A | 3.40 | 5.1 | FALSE |
| EP3A | 18.20 | 4.6 | FALSE |
| EP4A | 7.36 | 7.9 | FALSE |

### 2 Data analysis

We described the pipeline used in this analysis in the following repository: manuelsmendoza/elvira. There we posted the commands used to call the different tools required and the code created ad hoc for this analysis. We shared our code as an R package under Common Creative Licence. To describe a reproducible pipeline, we used a conda environment to do this analysis as described in the repository; the version of the different programs used are described in the environment information at the repository.

#### 2.1 Reads quality control

We checked the quality of the library using FastQC (Step 1). We detected two failures and two warnings. All of them are common in the quality assessment of RNA-Seq libraries (Figure S1).

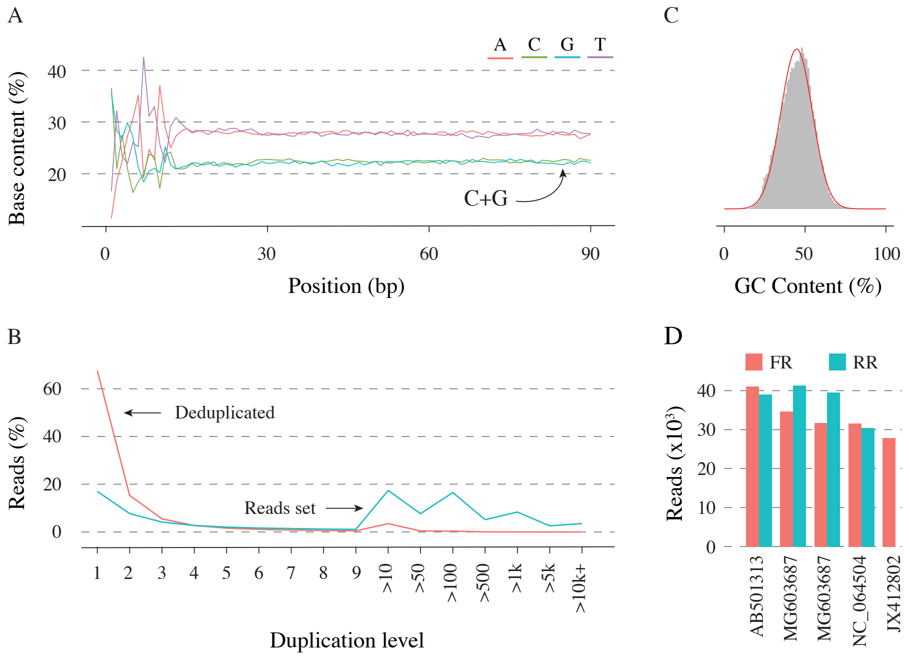

Figure S1: Modules that did not pass the quality assessment. (A) Base content per position. (B) Sequence duplication level. (C) Reads GC content distribution; the red shape represents the theoretical distribution. (D) Overrepresented sequences' abundance and clustered by their matches in Nucleotide NCBI database accession number (FR: forward reads; RR; reverse reads)

- Per base sequence content (FAIL): Nucleotides unbalance in the $15^{th}$ first position due to random hexamer priming (Hansen *et al.*, 2010).
- Sequence Duplication Levels (FAIL): Caused by the Differential transcripts expression (abundance) in the library as described in the documentation.
- Overrepresented sequences (WARNING): these sequences came from the remaining mitochondrial sequences fetched during library preparation (Smith, 2013). We detected five overrepresented sequences, four common in forward and reverse reads, matching all of

them with the small ribosomal RNA of different sea slugs (Table S2). We characterised the over-represented sequences by aligning them to the NCBI Nucleotide database using BLAST (blastn-short) (see repo). All the over-represented sequences matched the mitochondrial ribosomal small subunit. Based on this result, we tried to assemble the mitochondrial genome using different pipelines: MITGAR (Nachtigall *et al.*, 2021), Trimitomics (Plese *et al.*, 2019), GetOrganelle (Jin *et al.*, 2020) and NOVOPlasty (Dierckxsens *et al.*, 2017) but it was impossible to reconstruct the whole sequence, obtaining a set of non-contiguous transcripts up to 2k bp.

- Per sequence GC content (WARNING): caused by multiple options, e.g., the abundance of AT-rich regions from the mitogenome and also because the libraries of RNA-Seq present a high bias in the GC content between samples (Risso *et al.*, 2011).

We clipped sequencing adapters auto-detected by reads-pairs overlapping. After, we trimmed low-quality bases using the sliding-windows strategy (length 5 with reading mean quality 30). Finally, we corrected the miss-base calling (Step 2). The result of reads quality control is shown in Table S2.

Table S2: Reads quality control summary.

| | | |
|---|---|---|
| Reads passed filters | | 87.03% |
| Reads failed filters | Low quality | 12.74% |
| | Too short | 0.21% |
| | Low complexity | 0.01% |
| Estimated sequencing error | | 0.07% |
| Duplication rate | | 38.15% |

#### 2.2 Transcriptome assembly and annotation

We started the assembly by testing different combinations of parameters (Table S3) to obtain the best possible assembly. After each assembly, we evaluated them by attending to two parameters (Step 4):

- Completeness: Proportion of molluscan orthologs reconstructed (completely or fragmented).
- Correctness: Proportion of transcripts with a score greater or equal to the cutoff ("good transcripts").

We found a significant impact on the parameters tested in the final quality of the assembly. We used the combination of values that maximised the completeness and correctness of our transcriptome (Table S3). Despite this, we detected a poor reconstruction of molluscan orthologs in all the cases (less than 50%). Low biological quality may result from sequencing a highly fragmented (degraded) sample (Table S2).

Table S3: Result of testing different parameters to assemble the transcriptome of *E. viridis* using Trinity to get the optimal combination (pipeline Step 4).

| K-mer size | Min coverage | Min reads | Max diff | Correctness | Completeness |
|---|---|---|---|---|---|
| 20 | 1 | 2 | 2 | 0.85 | 0.33 |
| | | | 4 | 0.86 | 0.33 |
| | | | 16 | 0.85 | 0.33 |
| | | 10 | 2 | 0.88 | 0.31 |
| | | | 4 | 0.90 | 0.31 |
| | | | 16 | 0.96 | 0.31 |
| | 10 | 2 | 2 | 0.74 | 0.11 |
| | | | 4 | 0.77 | 0.11 |
| | | | 16 | 0.73 | 0.11 |
| | | 10 | 2 | 0.73 | 0.11 |
| | | | 4 | 0.72 | 0.11 |
| | | | 16 | 0.85 | 0.11 |
| 25 | 1 | 2 | 2 | 0.79 | 0.32 |
| | | | 4 | 0.84 | 0.32 |
| | | | 16 | 0.85 | 0.32 |
| | | 10 | 2 | 0.90 | 0.31 |
| | | | 4 | 0.90 | 0.31 |
| | | | 16 | 0.93 | 0.31 |
| | 10 | 2 | 2 | 0.70 | 0.10 |
| | | | 4 | 0.75 | 0.10 |
| | | | 16 | 0.77 | 0.10 |
| | | 10 | 2 | 0.78 | 0.10 |
| | | | 4 | 0.78 | 0.10 |
| | | | 16 | 0.83 | 0.10 |
| 30 | 1 | 2 | 2 | 0.76 | 0.29 |
| | | | 4 | 0.85 | 0.29 |
| | | | 16 | 0.86 | 0.29 |
| | | 10 | 2 | 0.87 | 0.28 |
| | | | 4 | 0.95 | 0.28 |
| | | | 16 | 0.91 | 0.28 |
| | 10 | 2 | 2 | 0.63 | 0.10 |
| | | | 4 | 0.74 | 0.10 |
| | | | 16 | 0.68 | 0.10 |
| | | 10 | 2 | 0.75 | 0.10 |
| | | | 4 | 0.78 | 0.10 |
| | | | 16 | 0.77 | 0.10 |

We obtained 41,072 raw transcripts even though only 16,912 were protein-coding sequences longer than 300bp (100aa). All the other transcripts may be another type of biological contaminants (Freedman *et al.*, 2021). In this project, we focused our analysis exclusively on the protein-coding sequences (CDS), removing all other sequences (Figure S2). Afterwards, we removed the redundant sequences (identical sequences present more than once). Lastly, we filtered the potential biological contamination from algae, plankton or any other source (pipeline step 6). We aligned the non-redundant transcriptome obtained previously (translated to proteins) to the NCBI Nr database and filtered the matches keeping only the transcripts that matched against a Tectipleura protein (Step 6) (Figure S2), the common taxa to the three genera *Elysia*, *Plakobranchus* and *Aplysia* (Kano *et al.*, 2016), the three genera analysed in this project. We used the Taxonomizr R package to obtain the taxonomic information from the protein matches.

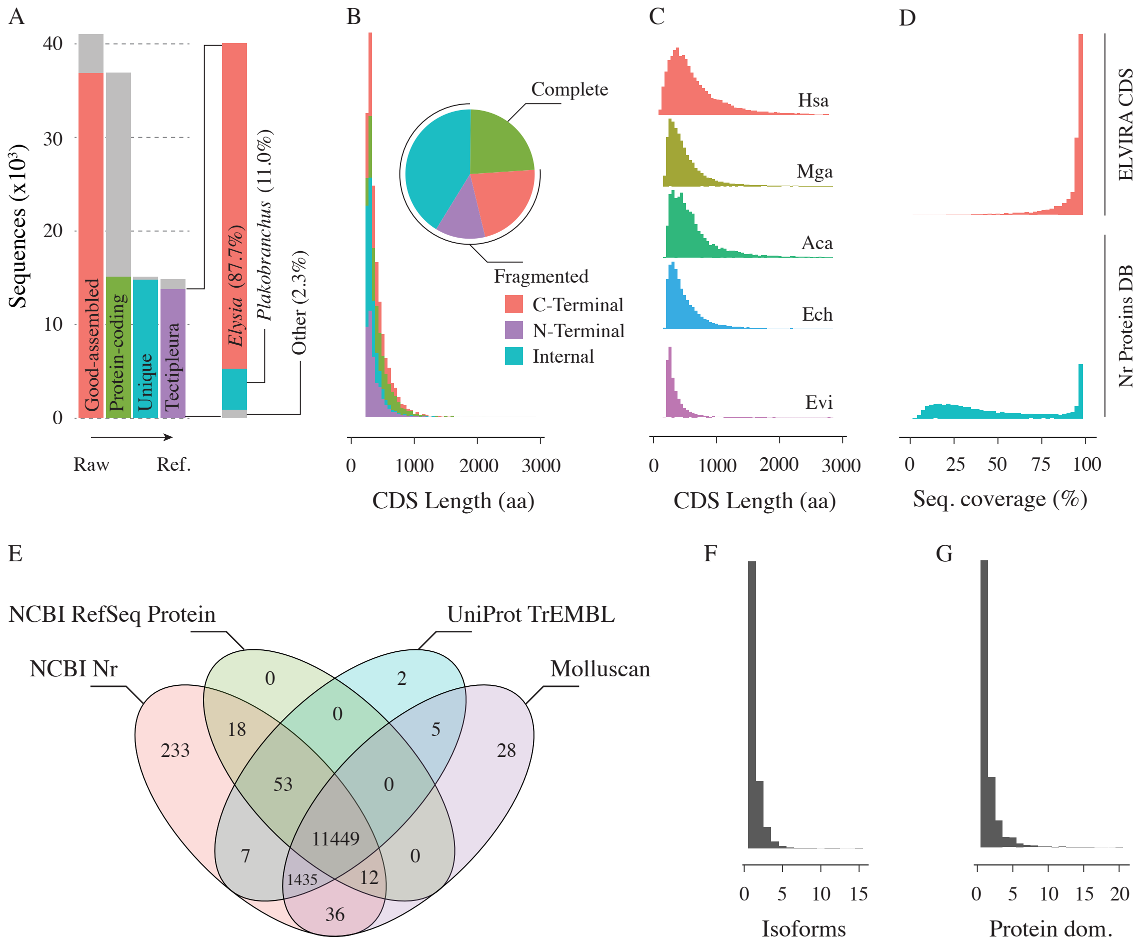

Figure S2: Transcriptome completeness evaluation and annotation. (A) In transcriptome filtering steps, the grey region represents the number of sequences discarded. (B) CDS length distribution, classified by completeness degree. (C) CDS length distribution comparison between different species; Hsa: *Homo sapiens*, Mga: *Mytilus galloprovincialis*, Aca: *Aplysia californica*, Ech: *Elysia chlorotica* and Evi: *Elysia viridis*. (D) Assembly CDS is covered by the DB proteins (upper), and proteins from the DB are covered by the transcriptome CDS (down). (E) Matches according to the four databases used; Molluscan: TrEMBL limited to molluscan entries exclusively. (F) Distribution the number of matches of the different proteins (with most having a unique match). (G) Protein domains were identified for CDS.

After applying the filters specified previously, we obtained a very fragmented transcriptome of reference (Figure S2 and Table S4). This reference transcriptome (ELVIRA, *Elysia viridis* reference assembly) comprised 12,884 CDS, from 261bp to 8,766bp. We annotated the transcriptome using a dual approach, firstly identifying the possible product of the transcript by homology (BLAST) (Altschul *et al.*, 1990; Camacho *et al.*, 2009; Shiryev *et al.*, 2007) (Step 7) and additionally identifying the protein domains of each CDS using HMMER (Johnson *et al.*, 2010; Eddy, 2011) (Step 9).

We used the results obtained using the UniProt TrEMBL DB to annotate the transcriptome because this database also contains information about the biological function of the proteins (Gene Ontologies, GO). We identified proteins and 3,538 protein domains from the Pfam DB (Fig. S2). 75% of the proteins identified had a unique match. However, there were also three proteins to which multiple CDSs matched: 15 matched to A0A3S1BCW4 (uncharacterised protein from *Elysia chlorotica* that contain two domains, coiled-coil and SH3), 12 matched to A0A433SR71 (Abhydrolase_5 domain-containing protein of *Elysia chlorotica*), and 11 matched to A0A3S1HW60 (Fibrinogen C-terminal domain-containing protein of *Elysia chlorotica*). The more abundant protein domains (more than 100 proteins) were WD40 (WD domain, G-beta repeat) and Pkinase (Protein kinase domain). 65% of CDS had a unique protein domain, and the maximum number of protein domains identified in the same CDS was 20, corresponding to ELVIRA04779. We annotated these CDS as EviOGT (*Elysia viridis* O-linked N-acetylglucosaminyltransferase), attending to the homology with A0A433SKQ2 and the protein domains (Glyco_transf_41, and several repetitions of multiple families of Tetratricopeptide repeat). Combining the protein domain information and the protein homology, we annotated the function of 9,333 CDS, 240 of them using exclusively the information about the protein domains, 5152 using only the protein homology information, and 3,941 mixing both methods.

Table S4: Assembly statistics from the raw assembly and the reference transcriptome.

| | **Raw** | **Reference** |
|---|---|---|
| N sequences | 41,072 | 12,884 |
| N bases (bp) | 32,118,314 | 9,301,944 |
| Smallesta (bp) | 301 | 261 |
| Largest (bp) | 18,577 | 8,766 |
| Mean length (bp) | 782 | 722 |
| N sequences $\leq$ 1kb | 8,477 | 2,457 |
| N sequences $\leq$ 10kb | 10 | 0 |
| N50 (bp) | 954 | 846 |
| GC (%) | 42 | 49 |
| Ort. Single-copy (%) | 25.8 | 26.1 |
| Ort. Duplicated (%) | 5.5 | 0.9 |
| Ort. Fragmented (%) | 3.7 | 3.4 |

Furthermore, we performed an additional annotation using eggNOG (Cantalapiedra *et al.*, 2021; Huerta-Cepas *et al.*, 2019) with the following options: Percentage identity 60%, Minimum % of query coverage 50, Minimum % of subject coverage 25, Gene Ontology evidence Transfer all annotations (including inferred from electronic annotation), PFAM refinement Realign queries to whole PFAM DB and SMART annotation activated.

#### 2.3 TSR identification and annotation

Thrombospondin type-1 repeat (TSR) proteins are involved in recognising hosts, pathogens, and symbionts (Adams, Tucker, 2000; Morahan *et al.*, 2009; Neubauer *et al.*, 2017). We manually annotated each transcript containing the protein domain of interest for each of the different TSRs, identifying 19 CDS that may belong to one of the five TSR families (Figure S3).

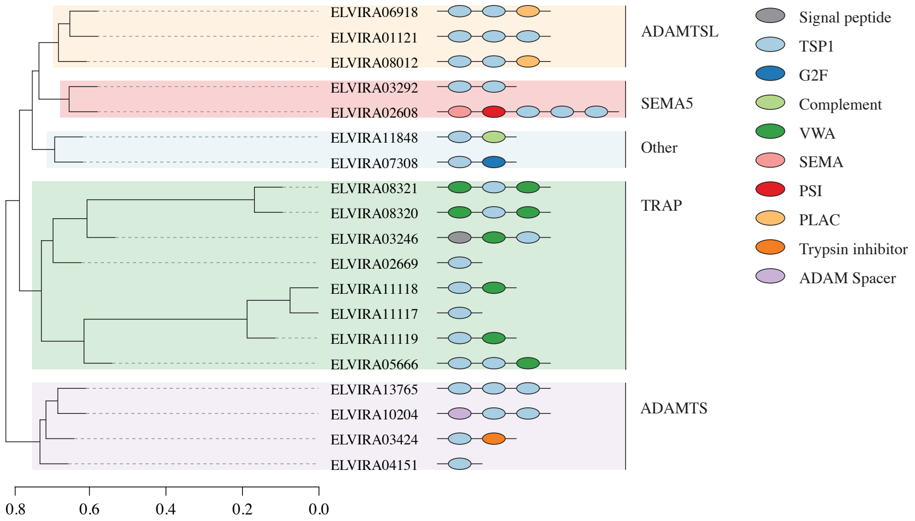

Figure S3: TSR superfamilies identified in the transcriptome of *E. viridis* clustered by protein identity. ADAMTS (a disintegrin and metalloproteinase with thrombospondin motifs); ADAMTSL (ADAMTS-like protein); SEMA5 (semaphorin 5B); TRAP (TNF Receptor Associated Protein), this group may include TRAP-like proteins too, but both were unified in a single group to use the same nomenclature that Neubauer *et al.*, 2017. Other correspond to the sequences we could not classify into a specific subfamily by lacking information.

Firstly, we filtered the CDS attending to the protein domains encoded after we re-ran homologies identification by local alignment (BLASTP) against the clustered Nr database and tried to identify the conserved domains architecture using CDART (Geer *et al.*, 2002) (we re-ran both steps at the NCBI BLAST Portal). Afterwards, we clustered the TSR-related CDS by sequence similarity to obtain the different subfamilies. We aligned the 19 CDS using MAFFT (Katoh *et al.*, 2002) and then estimated the sequence similarity using the R package seqinr. Finally, we clustered the sequences based on the similarity matrix. Furthermore, we also indenfied the transmembrane domains and signal peptides present in the CDS using DeepTMHMM (Hallgren *et al.*, 2022) and signalP (Teufel *et al.*, 2022), respectively.

We found two CDS that may encode a semaphorin 5 (SEMA5), one of them complete (ELVIRA02608) and the other fragmented (ELVIRA03292) (Figure S3). According to the standard description of the semaphorin families, the SEMA5 are not encoded by invertebrates (Pasterkamp, 2012), so we tried to infer if these CDSs are SEMA5 or any other protein with a similar structure (Figure S3). We did a phylogenetic analysis of SEMA5A, SEMA5B, and SEMA1 subtypes (Figure S4). We searched sequences of those families in the NCBI Protein database and fetched multiple sequences from diverse taxa. We aligned them using MAFFT and finally, we inferred an ML (maximum likelihood) tree using IQ-TREE 2 (Hoang *et al.*, 2018; Minh *et al.*, 2020). The evolutionary model was estimated using ModelFinder (Kalyaanamoorthy *et al.*, 2017). Despite detecting the SEMA5-like proteins in the transcriptome of *E. viridis* we did not find their receptors, the plexins (Plexin A1 and B3) (Nishide, Kumanogoh, 2018).

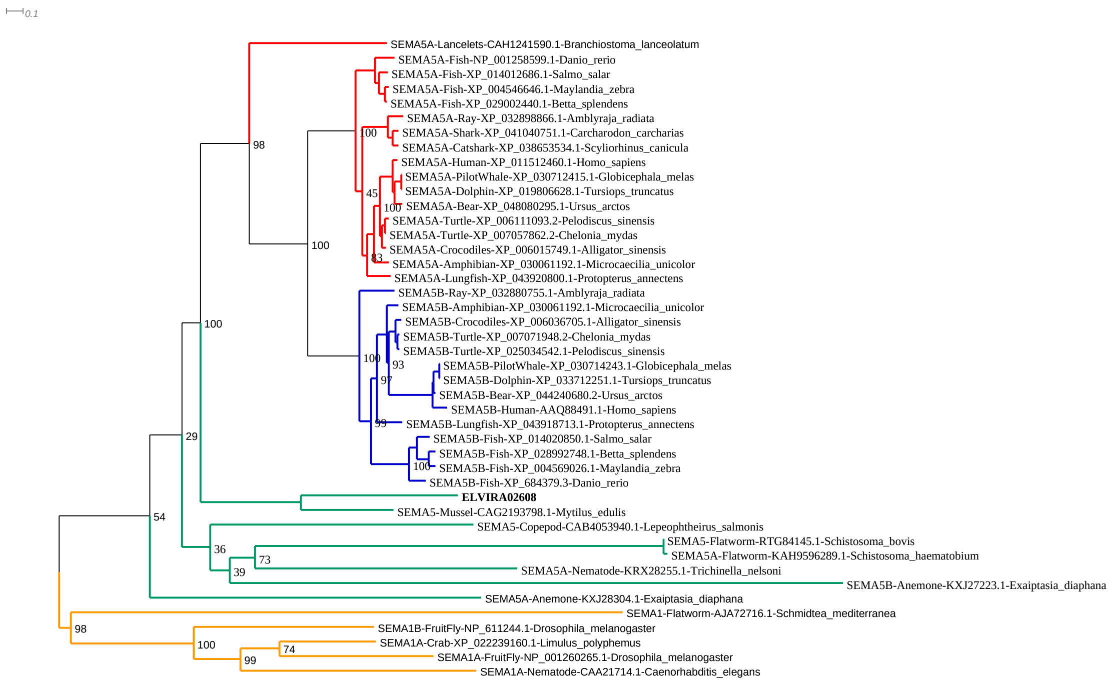

Figure S4: Maximum likelihood tree of three semaphorin families. We found two monophyletic groups corresponding to the vertebrates SEMA5B (blue) and the invertebrates SEMA1 (yellow); additionally, we found the vertebrates SEMA5A conform to a paraphyletic group (red), suggesting that both SEMA5 subtypes (A and B) diverged after the radiation of lancelets. The invertebrates' SEMA5 proteins (green) did not form a monophyletic group, instead defining multiple lineages.

#### 2.4 SR, TLR and CTLE identification and annotation

The method to identify the CDS that encode the different scavenger receptors (SR), Toll-like receptors (TLR) and C-type lectin receptors (CTLR) was the same described previously in the section related to TSR.

The obsolete kleptoplasts may be detected by the scavenger receptors (SR) that recognise oxidised lipoproteins (SR-B) and dead cells and debris (SR-B and SR-E) (Canton *et al.*, 2013). We could not reconstruct a complete SR even though we detected one or more scavenger receptor domains in 11 CDS (Figure S5). Most of them contained the SRCR domain exclusively, and none of the architectures detected corresponded to any of the standard structures of the different SRs.

Toll-like receptors (TLR) are membrane-bound pattern recognition receptors (PRR) that detect specific cell-surface components, e.g. triacylated lipoproteins (TLR2/TLR1), diacylated

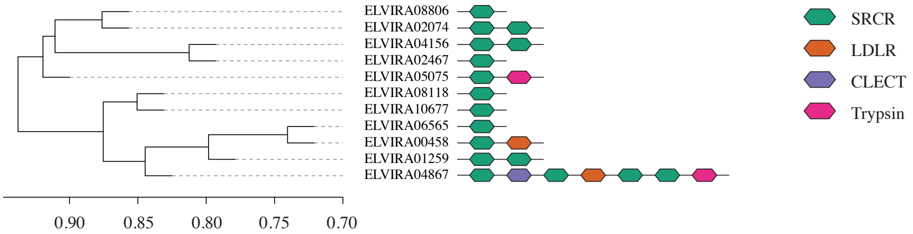

Figure S5: SR identified in the transcriptome of *E. viridis*. Scavenger receptor cysteine-rich domain (SRCR); LDLR (Low-density lipoproteins receptors); and CLECT (C-type lectin).

lipoproteins (TLR2/TLR6), lipopolysaccharides (TLR4) and nucleotides (TLR3, TLR7/TLR8 and TLR9) (Cao, 2016; Li, Wu, 2021). In molluscs, only two families have been described so far, TLR2 and TLR1 (Rauta *et al.*, 2014). We also detected the presence of TLR4 (Figure S6).

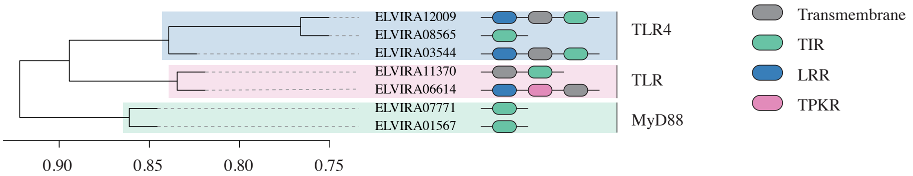

Figure S6: TLR identified in the transcriptome of *E. viridis*. Toll-interleukin receptor (TIR), Leucine-rich repeat domain (LRR), and TPKR (Tyrosine-protein kinase receptor). We assigned the two complete sequences and a fragment to TLR4 based on the protein homology. TLR group are fragmented CDS that exclusively have the extracellular domains (LRR) or the cytoplasmatic TIR domain. MyD88 are the CDS that have a single TIR domain without trans-membrane domains.

We identified 74 CDSs containing one or more C-type lectin domains (CTLD) (Fig. S7). The C-type lectins (CTL) are a very complex family of receptors that cover a wide range of ligands, e.g. proteins, lipids, and inorganic molecules (Mayer *et al.*, 2017). Possibly due to the high fragmentation of the assembly, none of these sequences had the classical structure composed of a trans-membrane domain followed by eight to ten CLTD and a head with a fibronectin type II domain together with a cysteine-rich domain (van der Zande *et al.*, 2021) (Figure S7). Yet, in sea slugs, the CTL-like proteins can present non-canonical domains, e.g. Astacin, Kringle and ShK (Melo Clavijo *et al.*, 2020).

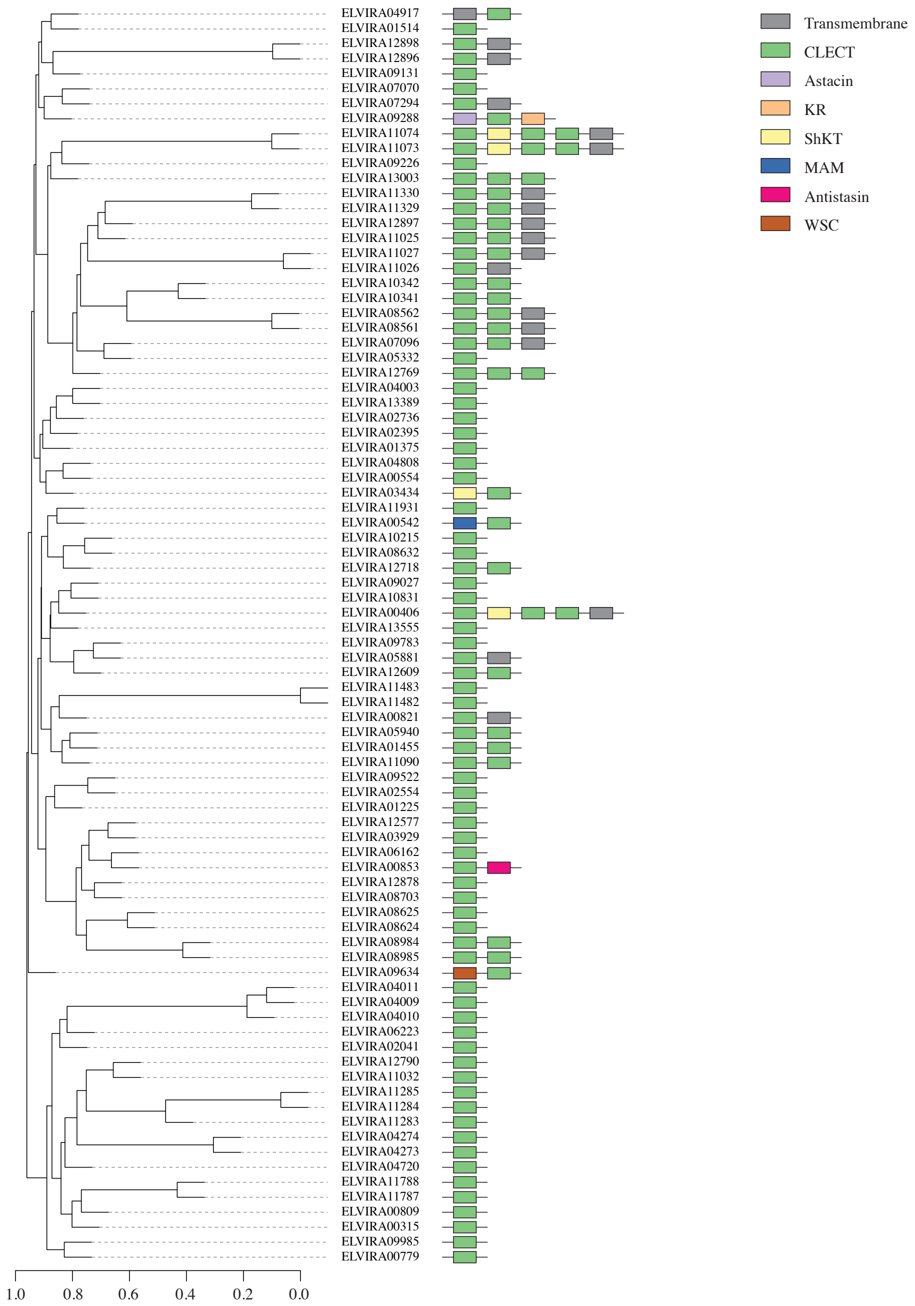

Figure S7: CTLR identified in the transcriptome of *E. viridis*. Scavenger receptor cysteine-rich domain (SRCR); LDLR (Low-density lipoproteins receptors); CLECT (C-type lectin).

### 3 Phylogeny

We tried to find the molecular mechanisms that allow some species to retain the plastids while the rest of the species can not do that (NR vs StrR and LtR). Furthermore, we also analysed the differences between the short- and long-term retainers (StrR vs LtR).

We assembled the transcriptomes of each species following the same steps used for the transcriptome of *E. viridis* (pipeline steps 1 to 6). The larger transcriptome corresponded to *E. crispata*, comprised of 106,224 sequences with 795bp of length on average. The most

Table S5: Information about the species used for the comparative analysis and their accession number in the NCBI SRA database.

| | **Source** | **Specie name** | **SRA Run** |
|---|---|---|---|
| | Carnivorous | *Aplysia californica* J. G. Cooper, 1863 | SRR6871219 |
| NR | Ulvophyceae | *Oxynoe viridis* (Pease, 1861) | SRR1505125 |
| StR | Ulvophyceae | *Elysia ornata* (Swainson, 1840) | DRR029459 |
| | | *Elysia cornigera* Nuttall, 1989 | SRR1582561 |
| LtR | Ulvophyceae | *Elysia timida* (Risso, 1818) | SRR1582571 |
| | | *Elysia crispata* Mörch, 1863 | SRR11015444 |
| | | *Plakobranchus ocellatus* van Hasselt, 1824 | DRR236388 |
| | Heterokonts | *Elysia chlorotica* Gould, 1870 | SRR8282417 |

complete transcriptome (attending to the number of molluscan orthologs assembled) was the *E. chloroticas*' assembly, containing 78.8% of orthologs. Only two samples (*E. chlorotica* and *E. timida*) had more than 75% molluscan orthologs. After removing possible contaminations and redundant sequences, the number of orthologs assembled remains equal (Figure S4 and Figure S6).

Table S6: Assembly statistics of the raw assemblies obtained for the different species. The molluscan orthologs are the sum of complete and fragmented molluscan orthologs detected in each transcriptome. The good CDS are the number of protein-coding sequences that passed the assembly cutoff of TransRate. The filtered sequences correspond to the final transcriptome after filtering by taxonomy and removing the redundant sequences.

| | **Transcripts** | **Transcripts Length (bp)** | | | Molluscan | Good | Filtered |
|---|---|---|---|---|---|---|---|
| | **assembled** | Max | Min | Mean | orthologs (%) | CDS | sequences |
| *A. californica* | 58,602 | 29,523 | 301 | 832 | 58.0 | 12,040 | 11,134 |
| *O. viridis* | 40,213 | 19,947 | 301 | 690 | 26.6 | 10,655 | 8,014 |
| *E. cornigera* | 67,816 | 22,511 | 298 | 916 | 66.0 | 27,760 | 18,895 |
| *E. ornata* | 90,962 | 14,963 | 298 | 879 | 60.4 | 30,000 | 19,692 |
| *E. chlorotica* | 86,649 | 26,662 | 282 | 960 | 78.8 | 30,960 | 20,099 |
| *E. crispata* | 106,224 | 12,786 | 301 | 795 | 49.7 | 24,150 | 16,001 |
| *E. timida* | 63,259 | 22,332 | 301 | 1039 | 77.5 | 23,915 | 17,323 |
| *P. ocellatus* | 76,675 | 26,086 | 301 | 936 | 69.4 | 22,648 | 15,866 |

Table S7: Number of orthogroups shared attending to different filters (same used in to obtain the transcriptome of *E. viridis*) and the number of CDS contained in those orthogroups from each species.

| | **Kleptoplasty** | **All LtR** | **Ulvophyceae-feeder LtR** |
|---|---|---|---|
| Total orthogroups | 573 | 4 | 3 |
| *A. californica* | 0 | 0 | 0 |
| *O. viridis* | 0 | 0 | 0 |
| *E. cornigera* | 940 | 0 | 0 |
| *E. ornata* | 878 | 0 | 0 |
| *E. chlorotica* | 908 | 7 | 0 |
| *E. crispata* | 902 | 7 | 3 |
| *E. timida* | 809 | 4 | 3 |
| *E. viridis* | 863 | 5 | 4 |
| *P. ocellatus* | 817 | 12 | 4 |

Table S8: Number of CDS and the total length of them used to reconstruct the phylogenetic tree.

| | **N CDS** | **N Bases** |
|---|---|---|
| *A. californica* | 1,160 | 373,090 |
| *O. viridis* | 1,258 | 328,286 |
| *E. cornigera* | 1,380 | 479,119 |
| *E. ornata* | 1,383 | 492,696 |
| *E. chlorotica* | 1,390 | 505,642 |
| *E. crispata* | 1,426 | 431,715 |
| *E. timida* | 1,376 | 483,352 |
| *E. viridis* | 1,299 | 368,376 |
| *P. ocellatus* | 1,373 | 480,934 |
